# Improved interpretability of bacterial genome-wide associations using gene cluster centric k-mers

**DOI:** 10.1101/2023.04.11.536385

**Authors:** Hannes Neubauer, Marco Galardini

## Abstract

The wide adoption of bacterial genome sequencing and encoding both core and accessory genome variation using k-mers has allowed bacterial genome wide association studies (GWAS) to identify genetic variants associated with relevant phenotypes such as those linked to infection. Significant limitations still remain as far as the interpretation of association results is concerned, which affects the wider adoption of GWAS methods on microbial data sets. We have developed a simple computational method (panfeed) that explicitly links each k-mer to their gene cluster at base resolution level, which allows us to avoid biases introduced by a global de Bruijn graph as well as more easily map and annotate associated variants. We tested panfeed on two independent data sets, correctly identifying previously characterized causal variants, which demonstrates the precision of the method, as well as its scalable performance. panfeed is a command line tool written in the python programming language and available at https://github.com/microbial-pangenomes-lab/panfeed.

## Introduction

Recent years have shown an unprecedented surge in bacterial genome sequencing. The availability of these large amounts of data has given rise to new opportunities in the field of microbial genomics, namely genomic epidemiology, population genetics and genome wide association studies (GWAS). For the latter the combination of ever larger sample sizes (*e*.*g*. > 200) and dedicated bioinformatic software has allowed the association of genetic variants to a number of bacterial phenotypes, some of which having high relevance to infection biology. These include virulence and pathogenicity^1^, antimicrobial resistance^2,3^, host range^4^ and carriage duration^5^. In many cases, statistically associated variants were verified to be causal for the phenotype through forward genetic techniques^1,6^, proving the power of the approach and its usefulness to elucidate crucial aspects of bacterial infections^7,8^.

While for human GWAS there are already a wide variety of computational approaches being used to run large scale analyses and for the interpretation of hits^9–11^, the field of bacterial GWAS cannot directly make use of these advancements because of several specificities in the genomic structure of microbes^7,8^. The clonal reproduction of bacteria leads to a stronger population structure in closely related strains. This in turn increases the linkage disequilibrium between genetic variants that contribute to a phenotype and those that do not, thus significantly reducing statistical power. Horizontal gene transfer (HGT) on the other hand increases the genomic plasticity of bacteria, leading to large differences in the genetic content across isolates. This makes the use of reference genomes, as is typically done in human-based studies, less appealing, since the large genomic heterogeneity cannot be captured by using a single reference genome; this is especially true for the extreme case of accessory genes not encoded in the chosen reference^12^. An approach that mitigates this reference bias is the use of pangenome graphs, which are often constructed using de Bruijn graphs (DBG) via k-mers^13^, unitigs^14^ or gene clusters^15^ as the smallest construction units. Pangenomes divide a species’ entire genomic information into two categories; the core genome, which contains sequences present within all or most (e.g. > 95%) strains of a species and an accessory genome that includes sequences occurring in a limited number of strains^16^. Thus far, k-mers and unitigs have proven to deliver a comprehensive set of causal variants, while providing a means to move away from single strain reference genomes. The drawback of such an approach is a lower interpretability of the resulting genetic variants, given that the local context of each variant is lost in the creation of the pangenome graph.

Some computational tools have been developed to partially overcome limitations in intepretability, namely DBGWAS^14^ and CALDERA^17^. DBGWAS (de Bruijn graph genome wide association study) for instance, constructs a compacted DBG (cDBG) and analyzes the resulting unitigs for association with the phenotype. As the original cDBG is quite large, DBGWAS outputs sub-graphs that focus on identified associations and their surroundings. CALDERA offers a similar approach to DBGWAS. It produces a cDBG and selects subgraphs to be tested for their associations to the phenotype, which reduces the computational workload. Both methods rely on the construction of a DBG, which scales poorly with the number of samples and is prone to artifacts due to the presence of repetitive regions in the input sequences.

Even though available bacterial GWAS methods have enabled discovery of variants related to many relevant phenotypes, interpretation and visualization of results tends to be cumbersome, as post GWAS processing of variants has often been implemented as an afterthought. Furthermore, the often used Manhattan plot only allows for visualization of variants that map to a specific reference, limiting interpretability of variants in the accessory genome of a species. While DBGs may give a good overview of the complexity of genomic diversity, they tend to still be difficult to interpret and scale poorly with increasing sample sizes.

To address these limitations we have developed a computational method (panfeed) to generate genetic variants from a collection of isolates to be used in a bacterial GWAS, with the main objective to improve the interpretability of the resulting associated variants. We leverage three key characteristics of bacterial genomes, namely the high density of coding regions and the empirical evidence that genes are often the main unit of molecular function and are transferred horizontally as a whole^18,19^. Encoding genetic variants using k-mers in the context of each gene cluster of the pangenome would therefore allow to account for both reference bias, since encoding variants using k-mers does not require the use of a reference, and for improved interpretability, since panfeed can retain the absolute and relative position of each k-mer in each input genome, thus allowing a fine grained mapping of associated variants. panfeed uses as input a genes’ presence/absence matrix (as the one given by Roary^20^, panaroo^21^ or ggCaller^15^) to create cluster specific k-mer sets and k-mer presence/absence patterns, which can be fed directly into bacterial GWAS software such as pyseer^22^. The resulting association table can easily be correlated to the gene clusters, which enables fast interpretation and visualization of variants occurring in both core and accessory genomes. This is made possible by the gene cluster centric approach of panfeed, which reduces the biases due to repetitive regions introduced by a “global” DBG. Furthermore, we show that resulting associations are comparable to those produced by using unitigs derived from a cDBG^14,23^, at a similar speed and memory consumption. Lastly, we show how a two-pass approach allows for faster processing with limited memory consumption and low disk space usage, which suggests that the approach can be scaled up to very large sample sizes.

## Methods

### panfeed algorithm

panfeed iterates over the gene presence/absence matrix produced by software such as Roary^20^, panaroo^21^ or ggCaller^15^, specifically the “gene_presence_absence.csv” file. For each gene cluster, the full nucleotide sequence of the gene is retrieved from the input GFF3 file of each of the input samples, optionally including sequences upstream and downstream of the start and stop codon, respectively. The canonical k-mers with a user-specified value of *k* (31 by default) are then generated, and a presence/absence vector across samples for each k-mer within the gene cluster is created, as well as the gene presence/absence vector. The algorithm also tracks the position of each k-mer in a user-provided list of strains, in absolute genomic coordinates and relative to the start codon of the gene. The data is then written into three separate plain-text tab separated files (TSV). The first file is “kmers.tsv”, containing the absolute and relative position of each generated k-mer for the samples indicated by the user. The second file is “kmers_to_hashes.tsv”, which contains the pairing between all generated k-mers and the identifier of their presence/absence pattern across all samples; we note that k-mers can share the same presence/absence pattern and that is therefore more computationally efficient to test each unique pattern rather than all k-mers. The last file is “hashes_to_patterns.tsv”, which contains a rectangular presence/absence binary matrix for each unique pattern; this file can directly be used by tools such as pyseer (option “--rtab”) to run a GWAS analysis. The first two files can be then used to directly map a set of patterns such as those passing the significance threshold to their precise location in the input samples. A visual representation of panfeed’s algorithm flow is shown in **Figure 1**.

**Figure 1:**
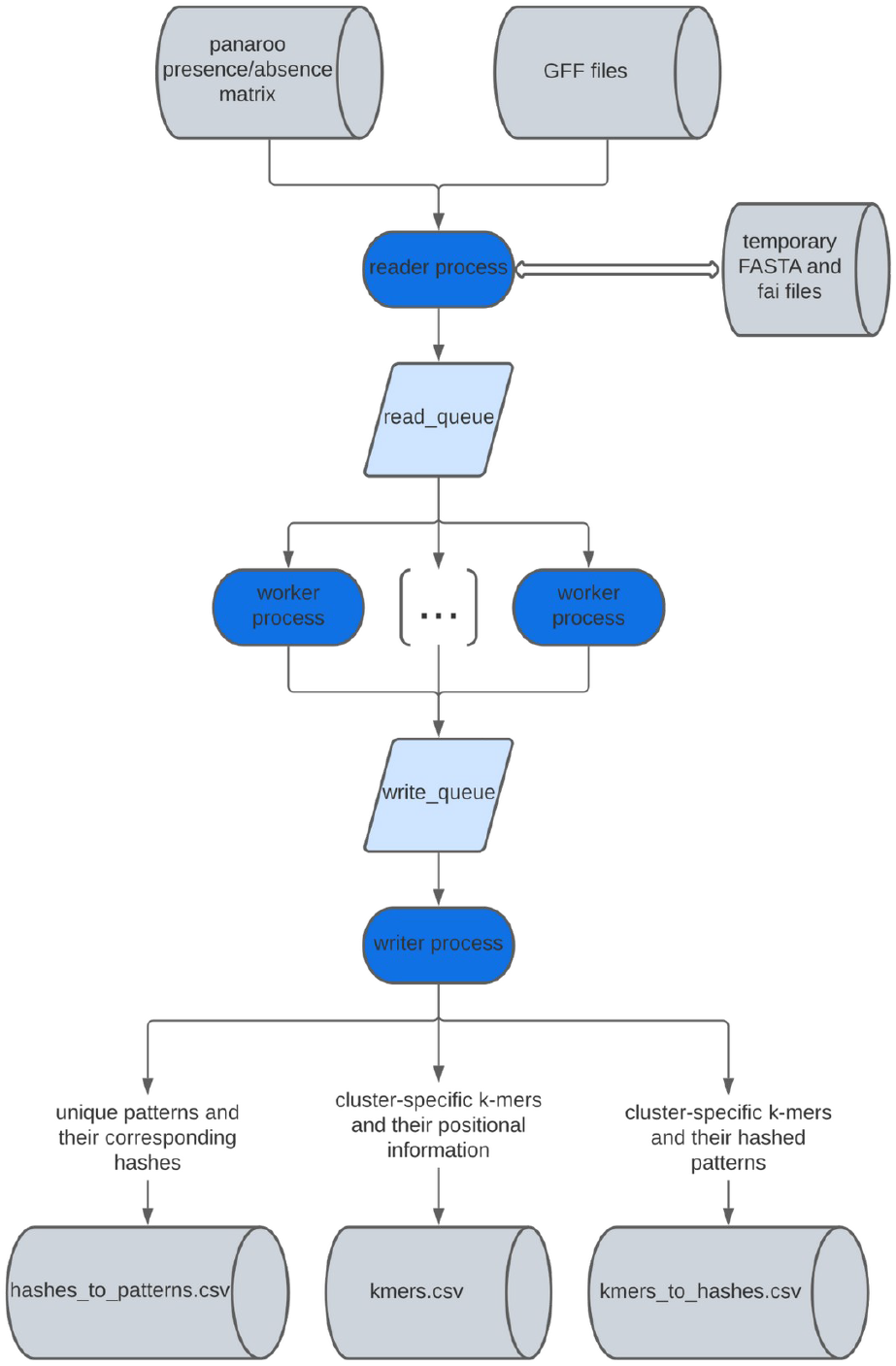
Structure of panfeed’s algorithm. Blue ovals represent sections of the algorithm that can be parallelized if the user provides at least 3 cores; the number of worker processes will be N-2, with N being the total number of provided cores.

panfeed can be sped up by taking advantage of the presence of multiple CPU cores; if at least 3 cores are provided by the user, then the algorithm is divided in 3 components that run in parallel, using N cores. The first one is the reader process, which cycles through each gene cluster and pushes the resulting sequences and gene coordinates in a queue, which is consumed by the second component, with N-2 separate workers, which extract the k-mers from each gene cluster and their coordinates and pushes them in a second queue. The last component is the writer process, which writes the three output files described above.

panfeed is written using the python programming languages and has the following dependencies, with the listed versions used in the benchmarks presented here: numpy v1.23.3^24^, pandas v1.5.0^25^, pyfaidx v0.7.1^26^, matplotlib v3.5.2^27^ and seaborn v0.11.2^28^. The code is available on github (https://github.com/microbial-pangenomes-lab/panfeed) and as a package on pypi (https://pypi.org/project/panfeed/) and on conda (https://anaconda.org/bioconda/panfeed).

### BSI data set

To evaluate the results of a GWAS using panfeed, both unitig-counter and panfeed were used to generate k-mer based genetic variants and fed to pyseer to find significant associations with the entry through the urinary tract for 912 *E. coli* strains causing bloodstream infections^29^ (BSI study). Resulting k-mers/unitigs were filtered using a Bonferroni correction threshold (alpha/family wise error rate 0.05) on the number of uniquely tested patterns. The overlap between gene cluster k-mers generated by panfeed and unitigs generated by unitig-counter was assessed with bedtools v2.30.0^30^ intersect, looking for overlaps of at least one base pair, disregarding the strandedness of sequences. The same command as in the original GWAS analysis by Denamur et al.^29^ was used to find associations with pyseer with both k-mers and unitigs.

### Neisseria data set

panfeed was also tested for its ability to interpret bacterial GWAS results on a data set of 1556 *N. meningitidis* strains^6^. Unlike the original study, we ran an analysis pooling the discovery and the replication data sets. **Figures 2B** and **3** were created using the convenience script provided with panfeed. Sequences for gene cluster “group_319” (*fHbp*) had to be manually shifted in a number of strains due to misannotation of start and stop codon positions in the GFF files (**Supplementary Figure 2**).

**Figure 2.**
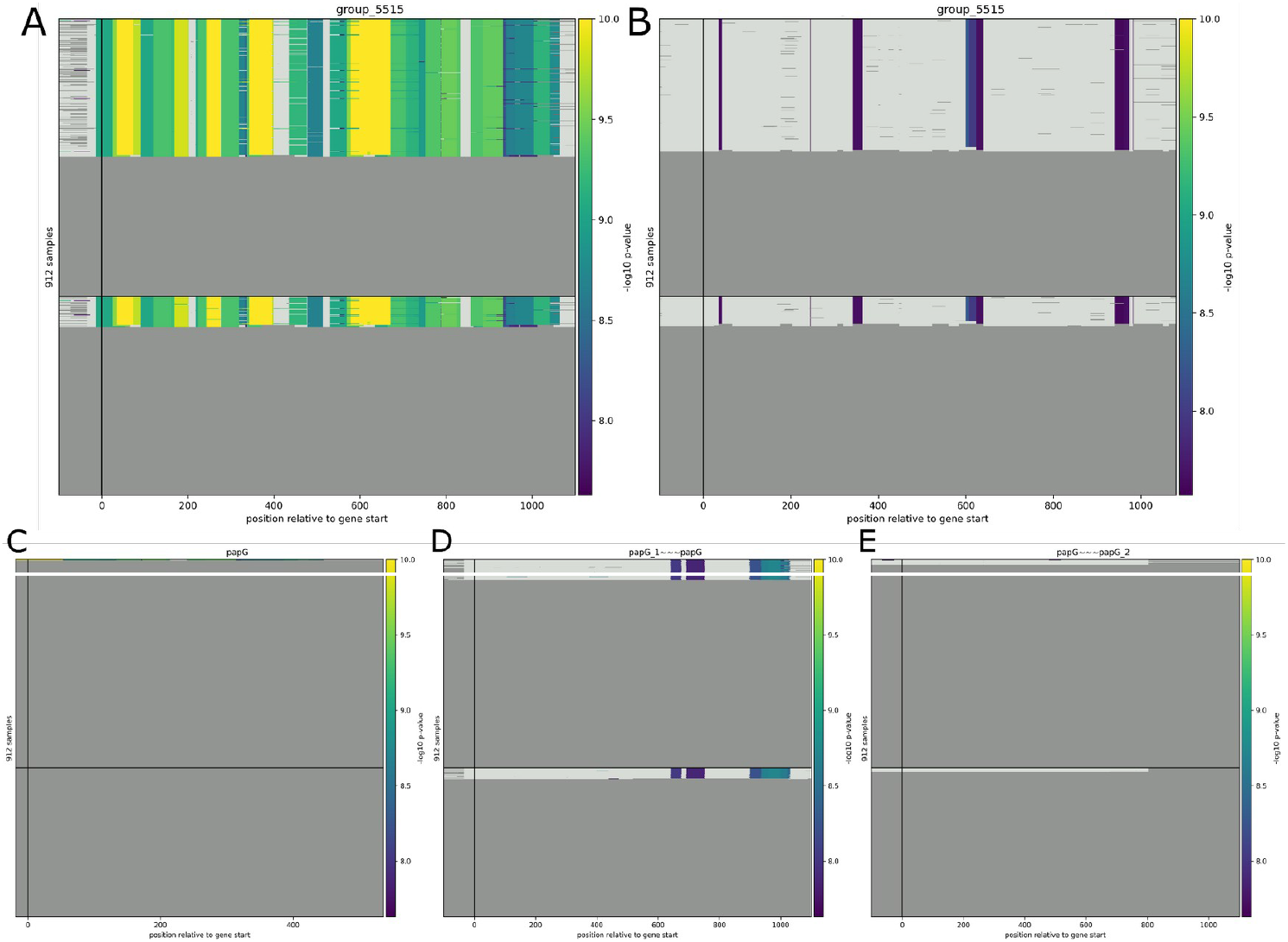
Association results comparison at single nucleotide resolution. Significantly associated unitigs (A, C, D, and E) / k-mers (B) are colored according to their -log10 p-value, ranging from the significance threshold up to 1E^-10^. Unitigs and k-mers whose association p-value is below the significance threshold are displayed in light grey. The X-axis represents the position relative to the gene start codon for each isolate’s genome, while each row represents a different isolate. The black vertical line displays the annotated gene start. The black horizontal line splits the graph between strains associated with urinary tract entry (top) and strains that are not (bottom). If a strain does not encode for the gene, the sequence will be displayed as dark gray. Panels A and B refer to gene cluster “group_5515”, panels (C), (D) and (E) display the significantly associated unitigs in clusters “papG”, “papG_1~~~papG”, and “papG~~~papG_2”, respectively.

### Benchmarks

The benchmarks for fsm-lite v1.0, unitig-counter v1.1.0^14,23^ and panfeed were conducted using the BSI (blood stream infection) study data set (see above). Subsets of 250, 450, 650 and the full 912 strains were created by randomly picking strains three times, meaning that every repetition contained a different strain composition, to minimize random effects. panfeed was set to record the k-mer positions for 237 strains, one for each of the lineages in the data set as determined by poppunk v2.4.0^31^. All benchmarks were performed on the same system, a workstation running Ubuntu 20.04.6 LTS with 36 Intel i9-10980XE cores, 128 GB RAM and a solid state disk (SSD 970 EVO Plus 2TB). The benchmarks for panfeed and unitig-counter were repeated with one core and four cores, while fsm-lite only supports the use of one core. The code used to benchmark panfeed, unitig-counter and fsm-lite can be found on github (https://github.com/microbial-pangenomes-lab/panfeed_benchmark_scripts).

We also tested an approach to optimize panfeed’s running time and memory consumption, which we term “two-pass approach”. The approach is implemented by giving panfeed no target strains to log k-mer positions for. This results in only the “hashes_to_patterns.tsv” and “kmers_to_hashes.tsv” files being populated, the former of which may be used as input for pyseer. Using panfeed’s convenience scripts, clusters containing significantly associated k-mers can afterwards be determined and used in the “--genes” option for a second pass of panfeed on the data set. This second run of panfeed subsequently logs the k-mer positions for the clusters given in the “--genes” option and the strains that were given as targets. A detailed scheme of the algorithm’s flow including the steps followed by the two-pass approach is available in **Supplementary Figure 1**.

## Results

### Gene cluster specific k-mers for fine grained mapping of significant associations

panfeed uses the gene-cluster centric pangenome creation of tools such as panaroo to reduce the complexity of the input strains’ pangenome, which removes the reference bias while allowing an easier interpretation of association results. Compared to DBG based approaches, panfeed avoids the creation of a global graph, which may introduce artifacts in the resulting presence/absence patterns of the genetic variants, such as those arising from repetitive sequences. panfeed extracts k-mers and their respective presence/absence patterns within each gene cluster, as well as the absolute coordinates and relative position with respect to the start codon. The resulting unique presence/absence patterns can be used for efficiently testing their association with a particular phenotype. Interpretability of associated patterns is aided by the fact that k-mers are already mapped to gene clusters, at base resolution level. Some example visualizations of the result of GWAS analyses enabled by panfeed are shown in **Figures 2** and **3**.

**Figure 3.**
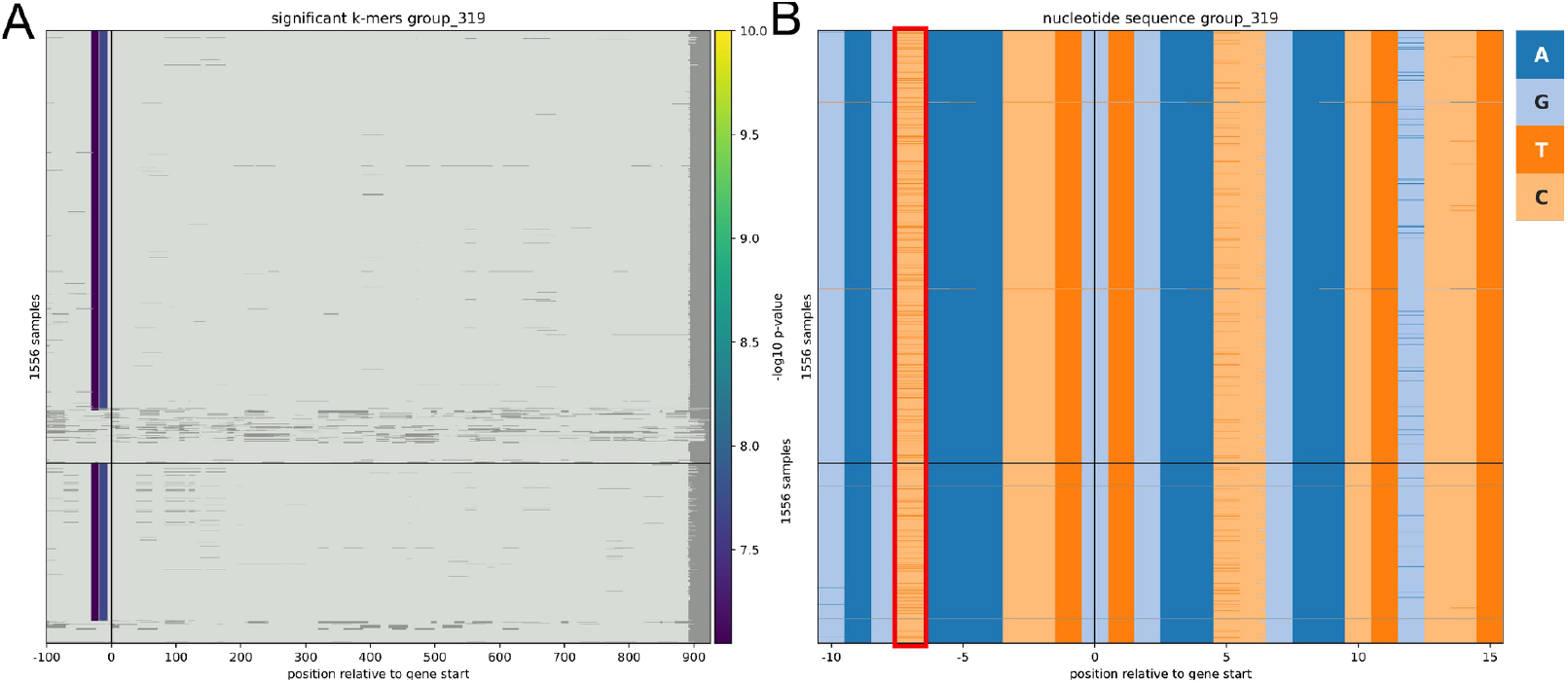
Fine mapping of association results. A) Significantly associated k-mers and their relative position with respect to the *fHbp* gene. Each row represents a different isolate. Some of the nucleotide sequences have been shifted due to a misannotated start codon. The black horizontal line separates IMD-associated strains (top) and carriage-associated strains (bottom). Dark grayareas are displayed, if a gene has a length below the longest gene within the specific cluster or the k-mers were filtered out due to having the same presence/absence pattern as the gene cluster B) Actual nucleotide sequences (encoded with different colors) across all isolates surrounding the *fHbp*_-7_T SNP, which is marked with a red box.

### Associations with gene cluster centric k-mers produce similar results as using global unitigs

To assess whether panfeed and unitig-counter, a DBG based method to produce joint core and accessory genome variation^14,23^, come to similar association results, we tested the overlap in significantly associated variants between unitigs output by unitig-counter and panfeed’s k-mers, using a previously published bacterial data set on *E. coli* bloodstream infections^29^ (BSI, see Methods). The data set contains 912 *E. coli* strains, for which the mode of entry into the host is available as well as confounding factors such as patient comorbidities. In total, we found 1,865,486 unique presence/absence patterns using panfeed and 2,132,621 using unitig-counter. Out of those we found a significant association with the target phenotype for 158,692 panfeed-generated k-mers and 29,728 unitigs. While the total numbers of significantly associated k-mers and unitigs differ, k-mers overlap with 79.9% of the unitigs and unitigs overlap with 89.6% of k-mers (**Table 1**), indicating a large agreement between the two variant sets. Overall we found these sequences to map to several gene clusters, 21 with global unitigs and 3 for panfeed’s k-mers (**Supplementary Table 1**). We found that the three gene clusters associated through panfeed were also found using global unitigs; the most prominent gene being *papGII* (named “group_5515” by panaroo’s automatic gene annotation), which encodes for an adhesin that is present at the tip of type P pili^32^. This adhesin has previously been linked to pyelonephritis and the urinary tract as portal of entry. The other two shared gene clusters were also fimbrial proteins such as *papF* (gene cluster “group_7980”) and *papD2*/*papD3* (gene cluster “papD_2~~~papD_3”), indicating how both approaches correctly identified the putative causal variant for entry through the urinary tract.

**Table 1:** Numbers of resulting unitigs and k-mers associated with entry through the urinary tract in the BSI data set and their overlap.

| Program | Total unitigs/k-mers | Overlapping unitigs/k-mers |
| --- | --- | --- |
| unitig-counter | 29'728 | 23'753 (79.9%) |
| panfeed | 158'692 | 142'180 (89.6%) |

We next leveraged panfeed’s ability to easily generate visualizations of the associations’ result to compare its performance with that obtained using global unitigs (**Figure 2**). Additionally, when using panfeed’s convenience scripts to connect k-mers and their association p-value, the presence of the entirety of gene cluster “group_5515” was flagged as significant, meaning that the k-mers derived from the *papGII* gene have associated with the phenotype with a low p-value. While this is not indicated in the visualization, it can be assessed from the table connecting k-mers, gene clusters and their respective association p-values. The association of *papGII* to the phenotype is in agreement with the hypothesis that the presence of the whole *papGII* cluster is causal for determining the entry through the urinary tract, rather than short variants in the gene^33,29^. The same association analysis using global unitigs resulted in similar areas of the gene being highlighted as highly significant as well as other regions being highlighted as less significant, while disregarding the possible association of the gene’s presence or absence with the phenotype of interest. In total, 67 significantly associated unitigs were present in gene cluster “group_5515”. We note that these significantly associated unitigs from cluster “group_5515” also occur in the clusters “papG”, “papG_1~~~papG”, and “papG~~~papG_2”, all annotated as *papGII*, albeit with different frequencies in the pangenome. This results in a difference in presence/absence patterns between k-mers and global unitigs, as panfeed’s k-mers are cluster-specific, meaning that the same k-mers in different clusters are only being considered as present in the cluster they occur in. This is in contrast to global unitigs, of which the presence or absence is being considered over the entire genome, allowing the same unitig occurring in different gene clusters to contribute to the overall presence/absence pattern, and thus potentially confounding the analysis.

### Simpler pangenome fine mapping of associated variants

The second data set we used to test panfeed consists of 1,556 *N. meningitidis* strains, for which phenotypic data on the induction of invasive meningococcal disease (IMD) in their hosts was measured^6^. Earle et al. found a SNP in the 5’ untranslated region of *fHbp* (*fHbp*_-7_T) that was significantly associated with IMD-positive strains in a discovery and replication data set consisting of a total of 1,556 strains. The GWAS analysis of Earle et al. found k-mers surrounding the SNP to have significant p-values, suggesting a causative link to the specific phenotype, which was then validated experimentally to influence the expression of *fHbp*, which in turn promotes immune evasion. As panfeed includes an option to incorporate promoter regions of genes in the association study, we also found significant k-mers spanning the same SNP, although with p-values just above the bonferroni-correction threshold. We then used panfeed’s convenience scripts to display the results of the association for this *fHbp* gene cluster and verify that it could correctly identify the experimentally verified SNP (**Figure 3**). The visualization allowed us to easily identify the genetic variant encoded by the k-mers as a SNP, given that the gene cluster is present in all strains and that the local context has a high level of homology across the isolates, which is not as straightforward when using global unitigs as the input genetic variants.

### panfeed has similar time and resource efficiency as DBG-based variant callers

We compared panfeed’s speed and memory consumption to fsm-lite and unitig-counter, which are programs commonly used to generate k-mers and unitigs for microbial GWAS. The benchmarks were performed using the 912 strain *E. coli* BSI data set^29^. To compare the programs’ performances at differently sized data sets, we generated four subsets with 250, 450, 650 and 912 strains. While both panfeed and unitig-counter have the option to be run with multiple cores, fsm-lite could only be run using one core. When running on one core, panfeed’s memory consumption is comparable to that of unitig-counter at 3.8 GB for 912 strains, while the running time for larger data sets exceeds that of fsm-lite at 3h 46min. Employing four cores enables panfeed to drastically reduce the running time to 2h 5min, while the memory consumption increases to a temporary maximum of 16GB as a trade-off (**Figure 4**). Users running panfeed on resource constrained systems should therefore use a single core or no more than three.

**Figure 4.**
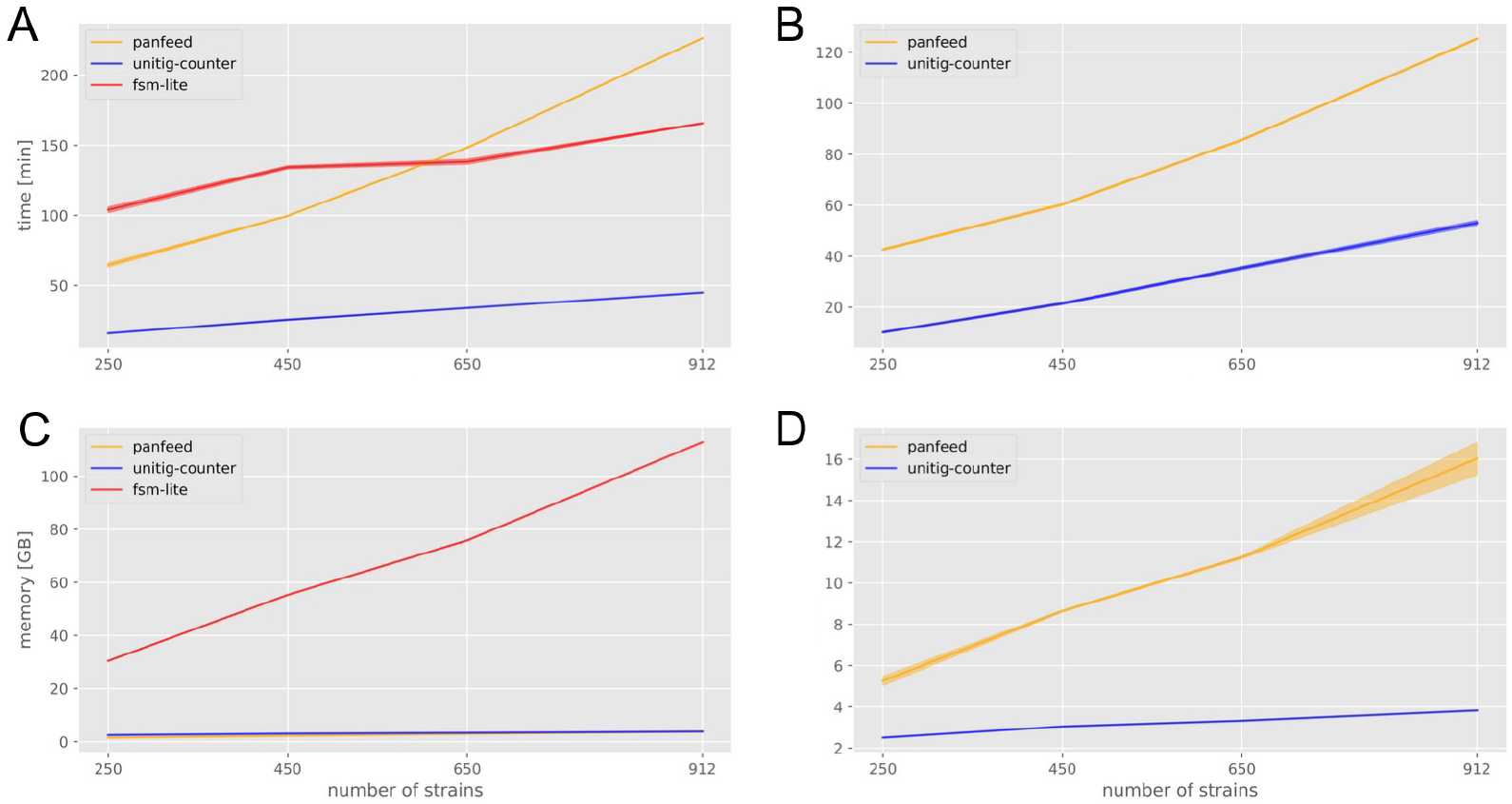
Time and memory consumption benchmarks of panfeed compared to fsm-lite and unitig-counter. Panels A and C were run using one core, while the benchmarks in panels B and D used four cores. Solid line represents the average over three replicates, shaded region the 95% confidence interval.

An alternative two-pass approach can be used, as described in the Methods section, to reduce both panfeed’s running time and memory consumption further. In this case, the first run of panfeed logs no k-mer positional information to file and is therefore used to generate the presence/absence matrix used as input for the associations. This first step on four cores takes 1h 18 min to finish and uses 14.7GB of peak memory. The second run, in which k-mers are logged only for those gene clusters that have at least one significant association, is completed in ~1 min with a peak memory consumption of 2.1 GB using one core. This two-pass approach additionally decreases the disk space that is needed to store the k-mer positional data from 500+ GB to 75.7 MB uncompressed, by only including k-mers from relevant gene clusters. The two-pass approach is therefore comparable in running time to the generation of unitigs, and likely able to scale up as data set sizes increase.

## Discussion

We have developed panfeed to address common difficulties in bacterial GWAS, using the principle that in bacteria a gene is the fundamental unit for many relevant phenotypes. This removes the necessity to map k-mers or unitigs back onto a reference genome, circumventing the drawbacks of using reference genomes. When comparing GWAS results of workflows using panfeed’s gene-centric k-mers versus DGB-derived unitigs, we were able to find most significant associations that were also found using global unitigs. The approach we developed does not suffer from biases that may be introduced from the construction of a DBG, but is obviously dependent on the approach used to annotate the input genomes and especially to cluster the resulting genes. An ideal data set would then have harmonized gene annotations across isolates. Recent advances in pangenome-aware gene calling and pangenome estimation might then be more appropriate; the recently described ggCaller^15^ algorithm’s output is in fact compatible with panfeed.

Mapping, annotation and visualization of k-mer based associations is a current limiting factor for wider adoption of bacterial GWAS. While encoding genetic variants as k-mers allows to encode both core and accessory genome variation, on the other hand the information on the nature and position of the genetic variant underlying a k-mer is not directly available. With panfeed however we have introduced ways to ameliorate these problems; each k-mer is explicitly mapped to gene clusters at single-base resolution, and both the nucleotide sequence and association statistics can be easily visualized via a heatmap. We have also shown how our approach also allows us to distinguish core and accessory genome variation by using the BSI^29^ and Earle et al.^6^ data sets. For the latter in particular we were able to find a localized association signal in the *fHbp* gene around the experimentally validated SNP *fHbp*_-7_, which was easily identified when displaying the nucleotide sequences at the positions with lowest association p-values. The capability of panfeed to incorporate promoter regions, in the context of their genes, in association studies, will likely help to illuminate a thus far underexplored field. We have therefore shown how panfeed can improve the current limitation on k-mer’s association interpretability, which will be important as data sets increase in size and complexity with the wider adoption of genome sequencing for applications in microbiology.

## Acknowledgements

HN and MG were supported by the Deutsche Forschungsgemeinschaft (DFG, German Research Foundation) under Germany’s Excellence Strategy - EXC 2155 - project number 390874280. HN was further supported by the DFG through grant GA 3191/1-1.

## Author contributions

HN and MG developed panfeed and its post-processing scripts. HN benchmarked panfeed and compared it to existing methods and prepared figures. HN and MG wrote the manuscript. All authors have no conflict of interest to report.

## Supplementary Material

**Supplementary Figure 1.**
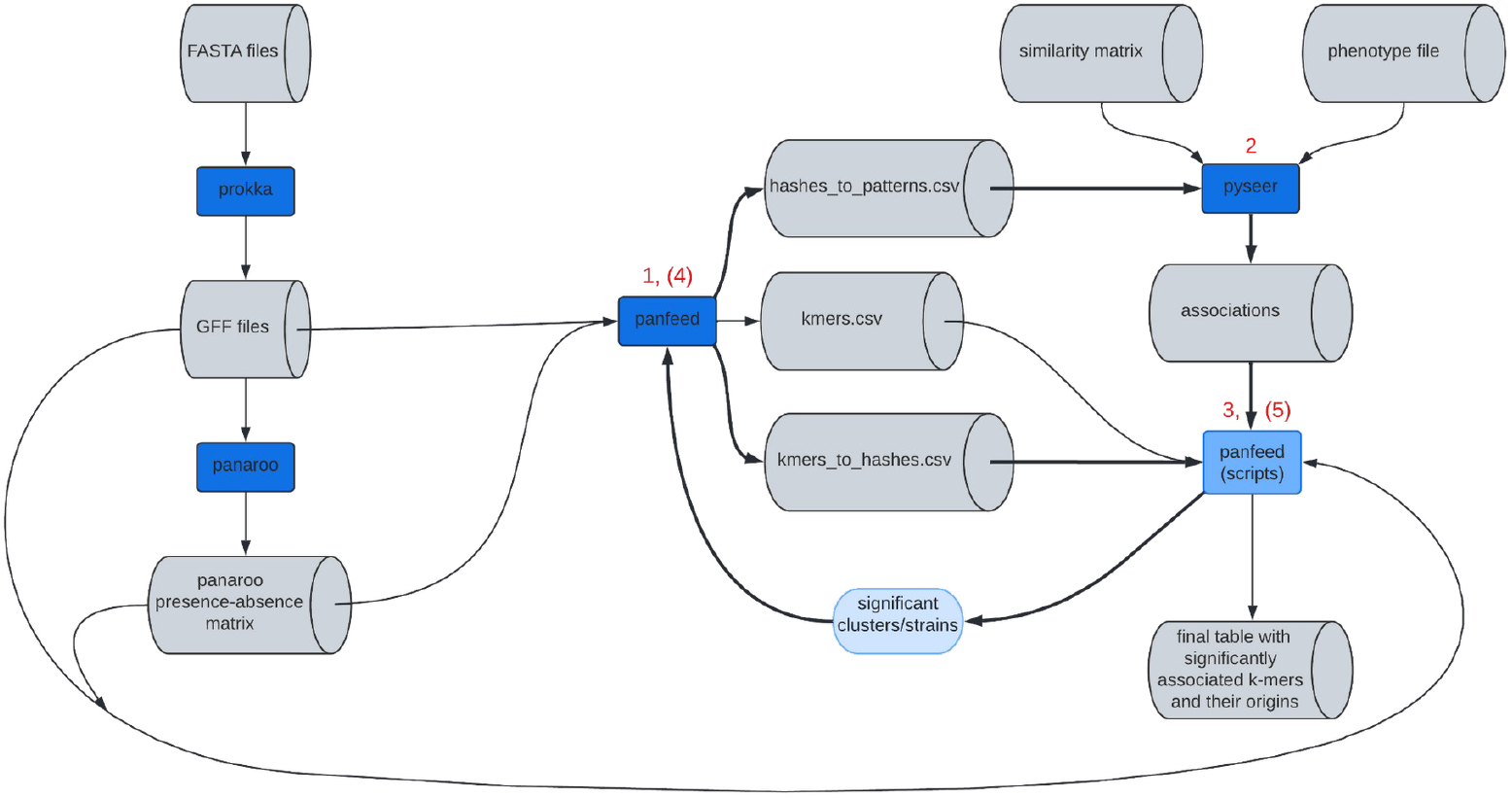
Workflow for a GWAS that employs panfeed as k-mer generator. Bold arrows mark steps that can be repeated to reduce both the memory footprint as well as resulting disk space being used by the written files. Red numbers indicate the step at which the specific program or script is being used. Steps in parentheses indicate optional steps for the two-pass approach that reduces time consumption and disk usage.

**Supplementary Figure 2.**
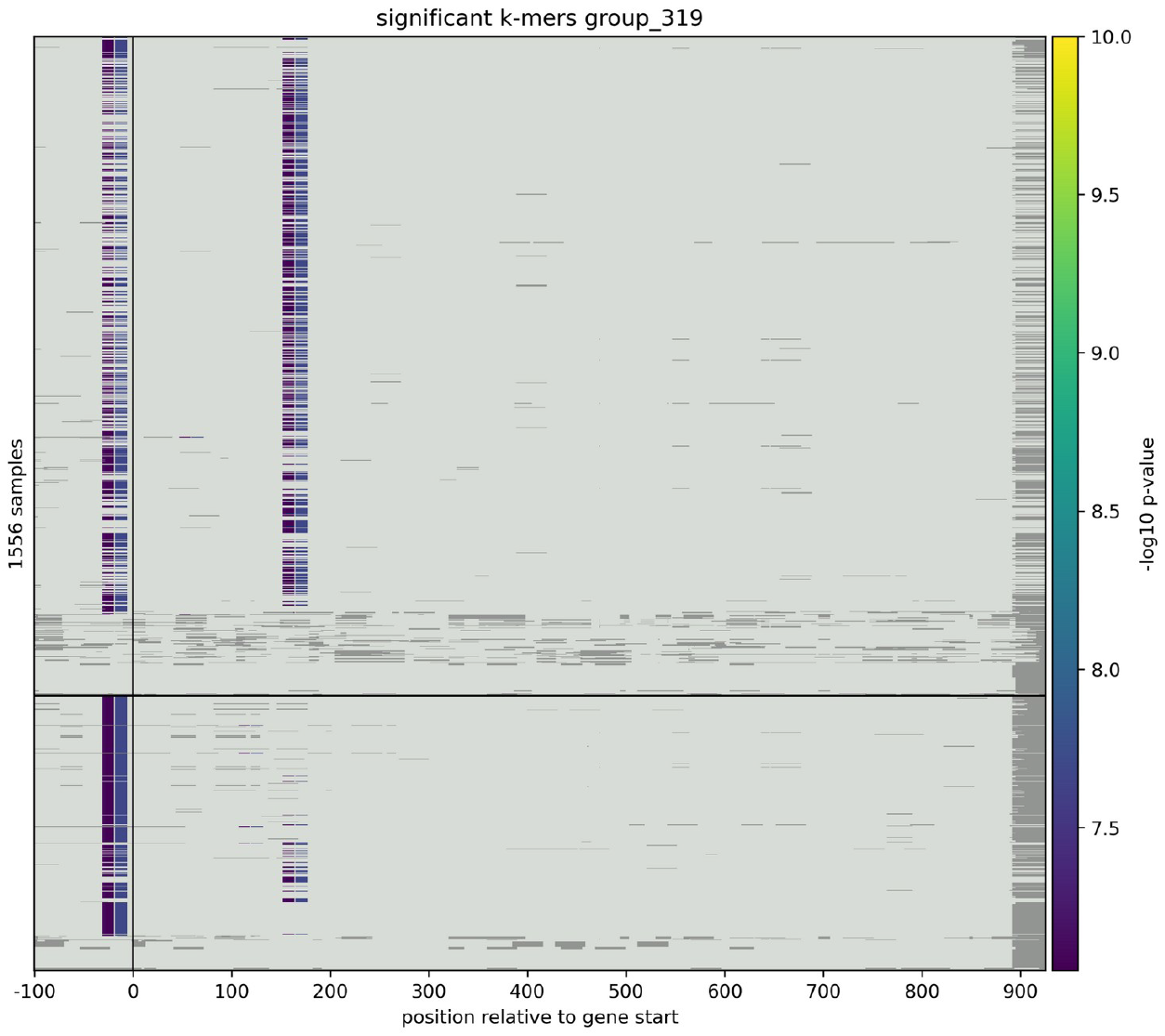
Unshifted -log10 p-values of significantly associated k-mers in gene cluster group_319 (*fHbp*) as found by panfeed in the *N. meningitidis* data set.

**Supplementary Table 1.** Gene clusters containing significantly associated unitigs/k-mers.

| Clusters with significantly associated unitigs | Clusters with significantly associated k-mers |
| --- | --- |
| group_11364<br>group_18381<br>group_19470<br>group_2782<br>group_5515<br>group_5516<br>group_6526<br>group_7266<br>group_7980,<br>intA_4~~~intS_2~~~intS_3~~~intA~~~intA_2~~~intA_3~~~intS_4~~~intA_1~~~int<br>A_5~~~intS_5~~~intS_1~~~intS<br>intA_7~~~intS_3~~~intA_6~~~intA_3~~~intA_1~~~intS_1~~~intA_5~~~intS_2~~~<br>intA_4~~~intS_4<br>intS_2~~~intS_4~~~intS_1~~~intA_5~~~intA_7~~~intA_1~~~intS_3~~~intA_4~~~<br>intA_6~~~intA_3~~~intS_6~~~intS<br>intS_3~~~intS_4~~~intA_2~~~intA_1~~~intS_1~~~intA_3~~~intA_7~~~intA_4~~~<br>intA_6<br>intS~~~intS_2, papC_4~~~papC_5~~~papC_2~~~papC_3~~~papC_1<br>papD_2~~~papD_3<br>papG<br>papG_1~~~papG<br>papG~~~papG_2<br>papH~~~papH_2<br>papK | group_5515<br>group_7980<br>papD_2~~~papD_3 |

